# Synaptic Dysfunction and Compensation After NMDA Receptor Ablation in the Mouse Medial Prefrontal Cortex

**DOI:** 10.1101/2025.08.14.669968

**Authors:** Rachel M. Dick, Lydia B. Cunitz, Aurora Torres Pérez, Habsa Ahmed, Anisha P. Adke, Cristina Rivera Quiles, Jason S. Mitchell, Ezequiel Marron Fernandez de Velasco, Nicola M. Grissom, Patrick E. Rothwell

**Affiliations:** Graduate Program in Neuroscience, University of Minnesota, Minneapolis, MN; Department of Neuroscience, University of Minnesota, Minneapolis, MN; University of Puerto Rico - Río Piedras, San Juan, PR; Medical Scientist Training Program, University of Minnesota, Minneapolis, MN; Department of Microbiology and Immunology, University of Minnesota, Minneapolis, MN; Department of Pharmacology, University of Minnesota, Minneapolis, MN; Department of Psychology, University of Minnesota, Minneapolis, MN

## Abstract

N-methyl-D-aspartate receptors (NMDARs) in the prefrontal cortex (PFC) are critical regulators of neuronal excitability, synaptic plasticity, and cognitive function. NMDAR disruptions, including pharmacological blockade and anti-NMDAR encephalitis, can mimic symptoms of schizophrenia. These observations support the glutamate hypothesis of schizophrenia, which posits that symptoms arise from abnormal corticolimbic glutamatergic signaling. Further evidence for this theory includes abnormal expression of NMDARs and decreased dendritic spine density in the PFC of individuals with schizophrenia, as well as altered spine density and synaptic transmission caused by genetic manipulation of NMDARs. However, it is unknown how progressive loss of NMDAR function in the PFC during adolescence – a developmental time period associated with significant synaptic pruning and symptom onset in schizophrenia – affects excitatory synaptic structure and function. In this study, we used in vivo genome editing to ablate expression of the Grin1 gene, which encodes the obligate GluN1 subunit of NMDARs, in medial PFC neurons of female and male adolescent mice. We assessed synaptic density and function in layer V pyramidal neurons at multiple time points using whole-cell patch-clamp electrophysiology, integrated with confocal imaging of dendritic spine architecture in recorded neurons. NMDAR ablation caused an early decrease in basilar dendritic spine density, followed by a rebound in spine density and corresponding increase in AMPAR-mediated synaptic transmission, suggesting that synaptic compensation maintains an allostatic set point. Our findings demonstrate that NMDAR ablation initially disrupts local PFC networks, followed by recovery via compensatory processes that could be impaired in disease states.

## INTRODUCTION

Abnormal glutamatergic signaling in the prefrontal cortex (PFC) is one of several biomarkers associated with schizophrenia [1]. Convergent evidence from genome-wide association studies [2,3], postmortem tissue histology [4,5], and in vivo neuroimaging [6,7] indicating altered expression and function of N-methyl-D-aspartate receptors (NMDARs). Early evidence for the link between glutamatergic signaling and schizophrenia came from studies showing that the administration of NMDAR antagonists such as phencyclidine (PCP) and ketamine induces a psychotomimetic state in healthy individuals [8,9] and worsen symptoms in patients [10–12]. Repeated NMDAR administration in rodents and primates also leads to cognitive deficits and progressive decreases in the synchronous firing of PFC neurons, which may be related to loss of synaptic connectivity [13–17]. This is consistent with human postmortem tissue studies demonstrating a reduction in dendritic spine density in patients with schizophrenia, specifically in the basilar dendrites of deep layer III neurons in the dorsolateral PFC [18,19]. These deficits in neural synchrony and spine architecture support the theory of activity-dependent disconnection [20–23], which posits that NMDAR loss drives spike timing defects in neurons and leads to subsequent weakening of PFC circuits [24].

NMDARs are heterotetrameric proteins containing two obligatory GluN1 subunits encoded by the Grin1 gene and two GluN2 and/or GluN3 subunits [25]. Although Grin1 mutations are not strongly associated with schizophrenia, manipulation of the Grin1 gene has served as a longstanding model for understanding the effects of decreased glutamatergic signaling. Initial research on the impact of NMDAR ablation upon neuroanatomy and physiology was conducted using mice with a homozygous genetic knockout of Grin1, which prevents the assembly of functional NMDARs [26]. However, *Grin1^−/−^* mice die 8-15 hours after birth due to respiratory failure, making them an unsuitable model for studying the long-term effects of NMDAR hypofunction [27]. The lethal effect of full Grin1 knockout can be avoided by using a more targeted approach; for instance, sparse Grin1 deletion in cultured cells or region-specific knockout in vivo. The impacts of these manipulations differ substantially depending on the brain region, developmental window, and method. Global knockdown of Grin1 to 10% of normal levels leads to reduced dendritic spine density in striatal medium spiny neurons [28] and unstable spines and reduced spine density in organotypic hippocampal slices [29]. In contrast, spiny stellate cells in layer IV of the barrel cortex have increased spine density in a cortical Grin1 knockout mouse [30].

These structural changes in dendritic spine architecture can also be associated with complex changes in excitatory synaptic transmission. Conditional genetic knockout of Grin1 in cortex and hippocampus with Nex-Cre mice leads to decreased spine density and a shortened lifespan of less than one month, but increased AMPAR-mediated synaptic transmission due to greater presynaptic axon bouton volume and postsynaptic density area [31]. Sparse conditional knockout of Grin1 in cultured hippocampal neurons also increases AMPAR-mediated synaptic transmission, but with no corresponding change in dendritic spine density [32]. This study also indicates that the timing of NMDAR manipulation may dictate functional consequences, as there was no change in AMPAR-mediated synaptic transmission when Grin1 was deleted in the hippocampus of mature mice (>P60). This temporal window for efficacy of NMDAR manipulation may vary between brain regions, as the effects of Grin1 deficiency on excitatory synaptic function in the mouse medial PFC (mPFC) can be rescued by restoring NMDAR expression in adulthood [33], indicating that the mature mPFC retains some of its capacity for neuroplasticity.

Taken together, the prior literature provides evidence that reduction of NMDAR function leads to changes in spine density and excitatory synaptic transmission, yet there is a critical lack of knowledge regarding the effects of NMDAR manipulation in the frontal cortex during adolescence. This is a relevant developmental period characterized by significant synaptic pruning [34] and the beginning of symptom onset in schizophrenia [35–39]. Adolescent manipulation of NMDAR function may also be relevant to anti-NMDAR encephalitis, which frequently develops during adolescence and produces symptoms that resemble acute psychosis [40,41]. To address these issues, we leveraged recent advances in genome editing that have made it possible to perform localized and chronic genetic manipulations in the mature nervous system in vivo [42,43]. We utilized clustered regularly interspaced short palindromic repeats (CRISPR)-Cas9 technology to delete the Grin1 gene encoding the obligate GluN1 subunit of NMDARs in mouse mPFC, and examined the effects of progressive loss of NMDAR function upon synaptic reorganization, by measuring synaptic strength and density at multiple time points during adolescence and early adulthood. We integrated electrophysiological measurements with confocal imaging of dendritic spines in order to understand how complex interrelated changes in neurotransmission and synaptic architecture may unfold over time in response to the loss of NMDARs.

## MATERIALS & METHODS

### Animals

Mice expressing a Cre-dependent Cas9-GFP transgene [44] were maintained on a C57Bl/6J genetic background. All experimental procedures were approved by the Institutional Animal Care and Use Committee at the University of Minnesota, and observed the NIH Guidelines for the Care and Use of Laboratory Animals. For additional details, see Supplementary Information.

### Surgical procedures

As previously described [45], mice between 5-6 weeks old underwent intracranial viral injections. Virus was infused bilaterally (500 nL each side at 120 nL/min) into the mPFC (AP: +1.80 mm, ML: ±0.35 mm, DV: −2.2 mm from bregma). Data were collected two weeks (14-27 days), four weeks (28-41 days), or six weeks (42-55 days) post-surgery. For additional details, see Supplementary Information.

### Immunohistochemistry for virus expression

Mice were deeply anesthetized using sodium pentobarbital (Fatal-Plus, Vortech Pharmaceuticals) and transcardially perfused with ice cold 0.01 M PBS followed by ice cold 4% PFA in 0.01 M PBS. Brains were extracted and post-fixed in 4% PFA at 4°C overnight, then transferred to a 10% sucrose solution. Following sectioning and immunohistochemistry, slices were mounted in PBS and coverslipped onto glass slides using ProLong Diamond Antifade mountant (Thermo Fisher Scientific). For additional details, see Supplementary Information.

### Brain slice electrophysiology

Coronal sections containing mPFC were collected as previously described [46]. Whole-cell voltage-clamp recordings were conducted in virally transduced pyramidal neurons in mPFC layer V. For additional details, see Supplementary Information.

### Confocal imaging and analysis of dendritic spines

Following whole-cell patch-clamp recordings with internal pipette solution containing neurobiotin, the pipette was slowly retracted until a gigaohm seal was formed, indicating successful separation from the membrane of the recorded cell that was preserved for morphological analysis. Dendritic spines were imaged using a Leica Stellaris 8 confocal microscope operated by a workstation running Leica LAS X acquisition software (version 4.6.1). All spine data was analyzed using Bitplane Imaris software (version 10.1). For additional details, see Supplementary Information.

### Statistical analysis

Similar numbers of male and female animals were used in all experiments, with sample sizes indicated in figure legends (n = number of cells, N = number of animals) and illustrated by individual data points in figures, with a visual distinction between cells from females (open circles) and males (filled circles). Sex was included as a variable in factorial ANOVA models analyzed using GraphPad Prism version 10, with repeated measures on within-subject factors. Significant ANOVA interactions were decomposed by analyzing simple effects (i.e., the effect of one variable at each level of the other variable). Significant main effects were analyzed using Fisher’s Least Significant Difference (LSD) post-hoc tests. The Type I error rate was set to α = 0.05 (two-tailed) for all comparisons. All summary data are displayed as mean +/− SEM. All figures were created in BioRender.

## RESULTS

### Validation of NMDAR Ablation using in vivo Genome Editing

To eliminate NMDAR expression, we used transgenic male and female mice expressing Cas9 and eGFP from the Rosa26 locus in a Cre-dependent fashion [44]. To activate Cas9 expression in mPFC neurons, we performed bilateral stereotaxic injection of an adeno-associated virus (AAV), with expression of mCherry-Cre driven by the human synapsin (hSyn) promoter sequence. The same AAV vector also expressed a guide RNA targeting the Grin1 gene [47], which encodes the obligate GluN1 subunit of NMDARs (Fig 1A). As a control, separate groups of mice were injected with a virus expressing a guide RNA for LacZ (a bacterial gene not present in mammals) in place of the Grin1 guide RNA. Stereotaxic injection of these AAV vectors was performed in adolescent mice at 5-6 weeks of age, a time point selected to mirror a developmental period associated with significant pruning of cortical networks [48,49] and the onset of schizophrenia-related symptoms in some patients [50]. We confirmed viral expression in the mPFC based on mCherry fluorescence, with co-localized GFP fluorescence by individual neurons indicating Cre-mediated transgene expression (Fig 1B).

**Figure 1.**
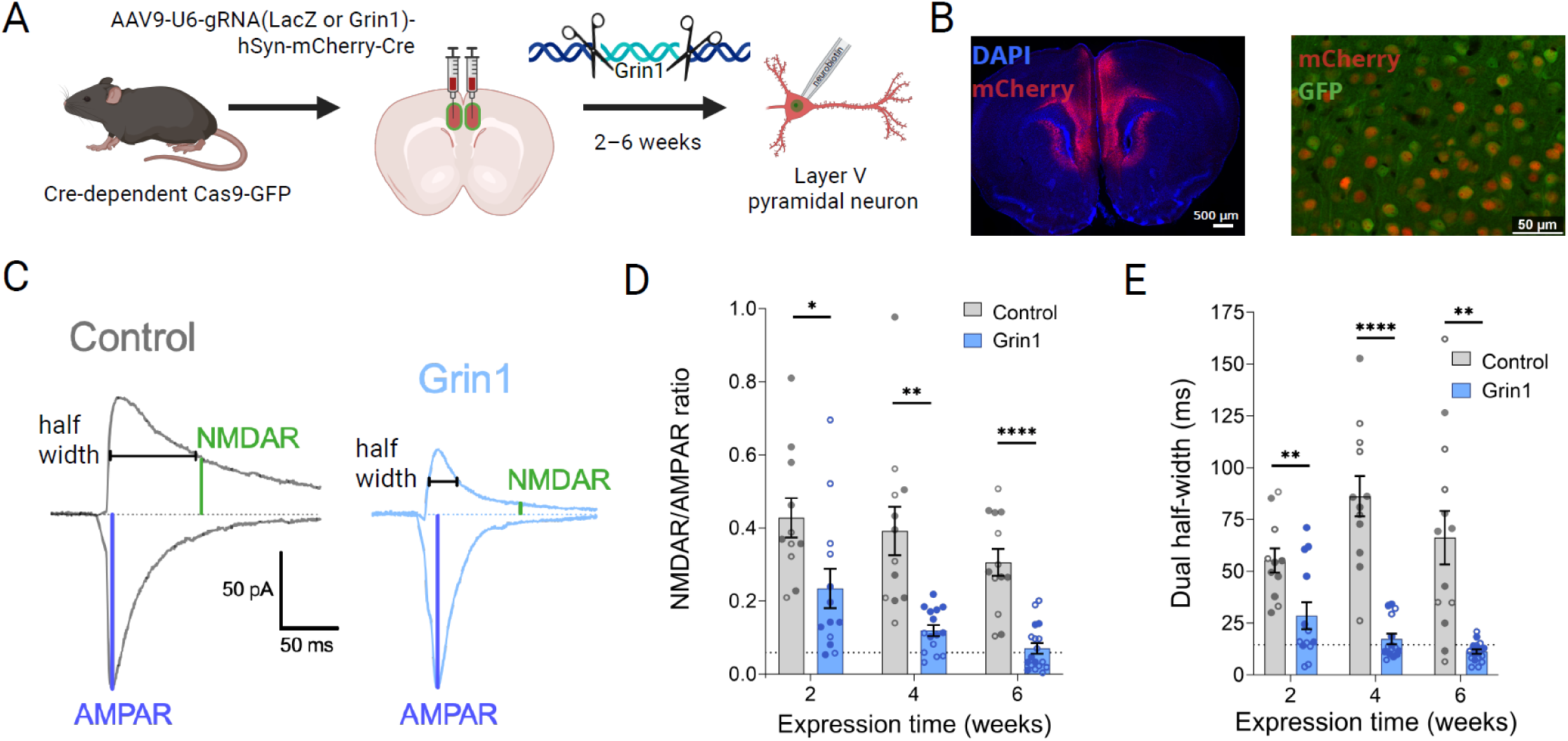
Grin1 ablation decreases NMDAR function. **(A)** Experimental overview showing mPFC intracranial injection of gRNA virus to ablate Grin1 gene expression, followed by whole-cell patch-clamp recording and neurobiotin cell filling. **(B)** mPFC expression of mCherry and GFP in virally transduced cells (left: 10x magnification tiled image, right: 20x magnification). **(C)** Sample traces of evoked excitatory postsynaptic currents (EPSCs) at −70 mV and +40 mV to measure AMPAR and NMDAR currents, respectively. **(D)** Grin1 ablation reduces NMDAR/AMPAR ratio with a significant main effect of virus (F_1,70_ = 58.80, P < 0.0001) (control n/N=11/4, 12/5, and 13/6 at 2, 4 and 6 weeks; Grin1 n/N=13/4, 15/6, and 18/6 at 2, 4 and 6 weeks) and **(E)** half-width of the dual component EPSC with a significant main effect of virus (F_1,70_ = 74.74, P < 0.0001) and time x virus interaction (F_2,70_ = 3.670, P = 0.0305) (same sample sizes as D). Dotted lines indicate values for NMDAR/AMPAR ratio (D) and dual half-width (E) measured under pharmacological blockade of NMDAR function with 50 µM D-APV (n=4). Data are mean ± s.e.m. for all panels; open and closed circles indicate recordings from female and male mice, respectively. *P < 0.05, **P < 0.01, ****P < 0.0001; three-way ANOVA followed by simple effect test; see Data S1 for complete statistics. https://BioRender.com/8iil7j5

To confirm successful ablation of the Grin1 gene and subsequent loss of NMDAR function in virally transduced cells, we performed whole-cell patch-clamp recordings from layer V pyramidal neurons in acute mPFC brain slices, prepared 2-6 weeks after stereotaxic surgery. NMDAR current was measured by recording electrically-evoked excitatory postsynaptic currents (EPSCs) at +40 mV and calculating the amplitude at 50 ms post-stimulus, by which point much of the fast AMPAR current has decayed (Fig. 1C) [51]. AMPAR current was measured by recording EPSCs at −70 mV and calculating the peak amplitude. At all timepoints, there was a significant reduction in the NMDAR/AMPAR ratio in the Grin1 group compared to the control group (Fig 1D). The half-width of the dual component EPSC (NMDAR+AMPAR) recorded at +40 mV was similarly reduced at all timepoints (Fig 1E). This indicates that the change in the NMDAR/AMPAR ratio is due to a loss of NMDARs, which have slower kinetics and therefore produce a more sustained excitatory current at +40 mV compared to AMPARs. After six weeks of Grin1 virus expression, decreases in average NMDAR/AMPAR ratio and dual EPSC half-width were comparable to pharmacological blockade of NMDAR function with 50 µM D-APV (dotted line in Fig 1D-E), indicating robust loss of NMDAR function.

### Dendritic Spine Architecture after NMDAR Ablation

To quantify synaptic density after ablation of NMDAR expression, we filled individual neurons with neurobiotin during whole-cell patch-clamp recordings, and then performed confocal imaging of dendritic architecture in the same pyramidal neurons (Fig 2A). Using Imaris software, spines were also automatically classified into four morphological groups: mushroom spines, which tend to be more stable and associated with greater synaptic strength; and filopodia/dendrites, stubby spines, and long/thin spines, which are more immature and weaker [52].

**Figure 2.**
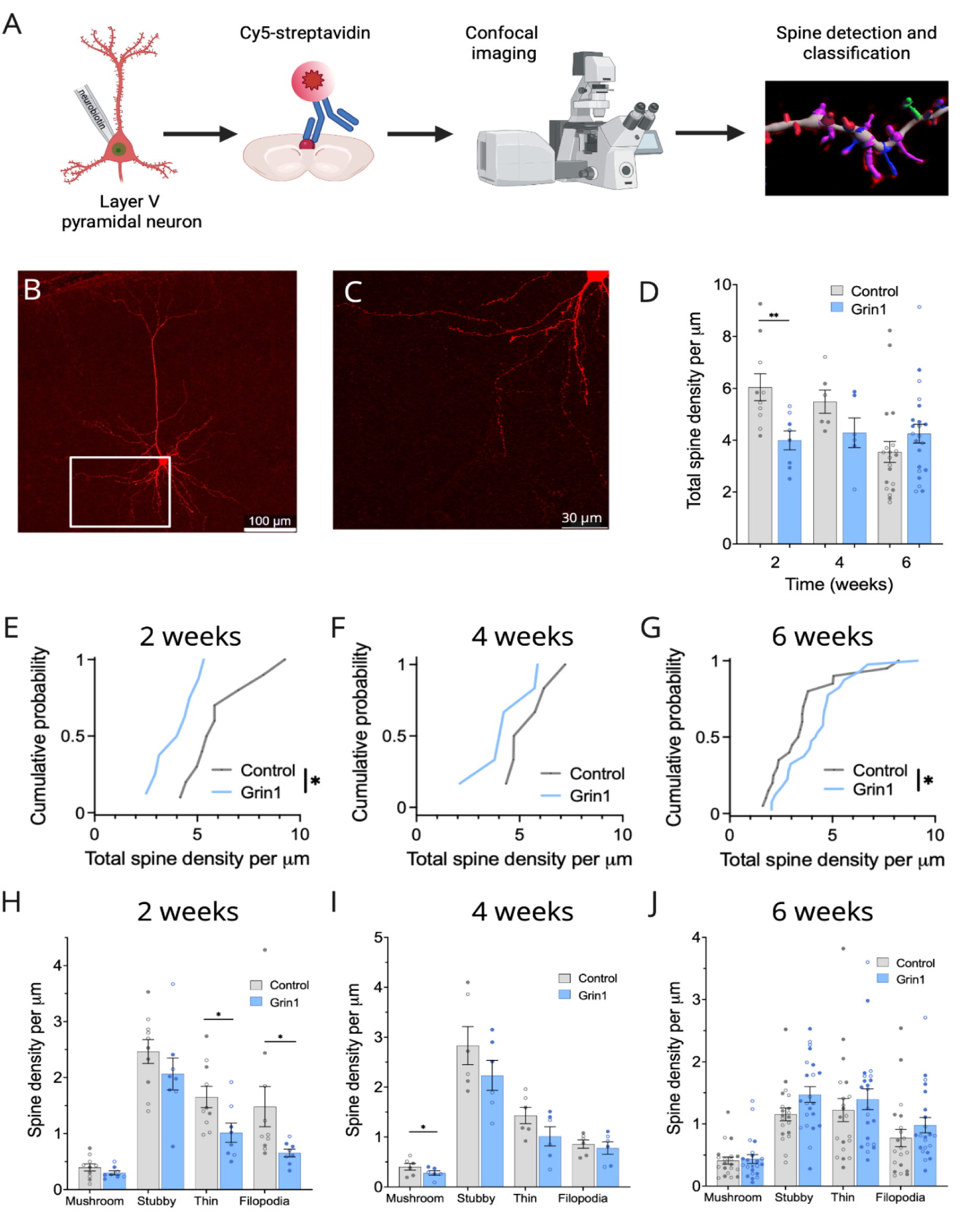
NMDAR ablation causes an initial decrease and subsequent increase in basilar spine density. **(A)** Schematic showing neurobiotin filling, imaging, and analysis of dendritic spines. **(B-C)** Sample image of the basilar dendrites of a layer V mPFC pyramidal neuron at 20x magnification (B) and 63x magnification (C). **(D)** Decrease in total basilar spine density in the Grin1 group at two weeks, with a trend towards increased density at six weeks (main effect of virus, F_1,60_ = 4.709, P = 0.0340; virus x time interaction, F_2,60_ = 5.307, P = 0.0075) (control n/N = 10/5, 6/4, and 20/12 at 2, 4 and 6 weeks; Grin1 n/N = 8/4, 6/4, and 22/8 at 2, 4 and 6 weeks). **(E, H)** Basilar spine density distribution is significantly reduced in the Grin1 group at two weeks (D = 0.6000, P = 0.0499), driven by significant reductions in thin spines (F_1,14_ = 5.971, P = 0.0284) and filopodia (F_1,14_ = 4.935, P = 0.0433). **(F, I)** Density of mushroom spines is significantly reduced at four weeks (F_1,8_ = 6.689, P = 0.0323). **(G, J)** Basilar spine density distribution is significantly increased at six weeks (D = 0.4364, P = 0.0370), with a trend towards increased density in immature spine categories. Sample sizes for panels E-J are the same as panel D. Data are mean ± s.e.m. for all panels; open and closed circles indicate recordings from female and male mice, respectively. *P < 0.05, **P < 0.01; three-way ANOVA followed by simple effect test (D); Kolmogorov-Smirnov test (E-G); or two-way ANOVA (H-J); see Data S1 for complete statistics. https://BioRender.com/46hxi18

In the basilar dendritic arbor (Fig 2B-C), we observed differences between the Grin1 and control groups that varied across time. At 2 weeks, there is a significant decrease in basilar spines in the Grin1 group (Fig 2D), with a corresponding leftward shift in the cumulative probability distribution (Fig 2E), driven primarily by a loss of thin spines and filopodia (Fig 2H). This same trend was observed to a lesser extent at 4 weeks (Fig 2F), with a significant decrease in the densities of mushroom spines (Fig. 2I). At six weeks, this pattern reversed (Fig. 2D) and the cumulative probability distribution shifted rightward toward a significant *increase* in the Grin1 group compared to LacZ (Fig 2G), a pattern most apparent in the less mature spine categories (Fig. 2J). These morphological changes were specifically observed in the basilar dendritic arbor, as no significant changes in total spine density or specific spine categories were observed at any time point in the apical dendritic arbor of the same cells (Fig. S1).

### Functional Correlates of Synaptic Strength and Density

To investigate the functional implications of these alterations in synaptic architecture, we recorded miniature excitatory postsynaptic currents (mEPSCs) in the presence of tetrodotoxin (500 nM), providing a measure of spontaneous excitatory neurotransmission (Fig 3A). The mEPSC amplitude, which is a correlate of postsynaptic strength, was not significantly changed at any time point (Fig 3B). However, there was a significant increase in mEPSC frequency at 6 weeks (Fig 3C), with a corresponding leftward shift in the cumulative probability distribution of mEPSC inter-event intervals at 4 and 6 weeks (Fig 3F and 3H), but not at 2 weeks (Fig 3D).

**Figure 3.**
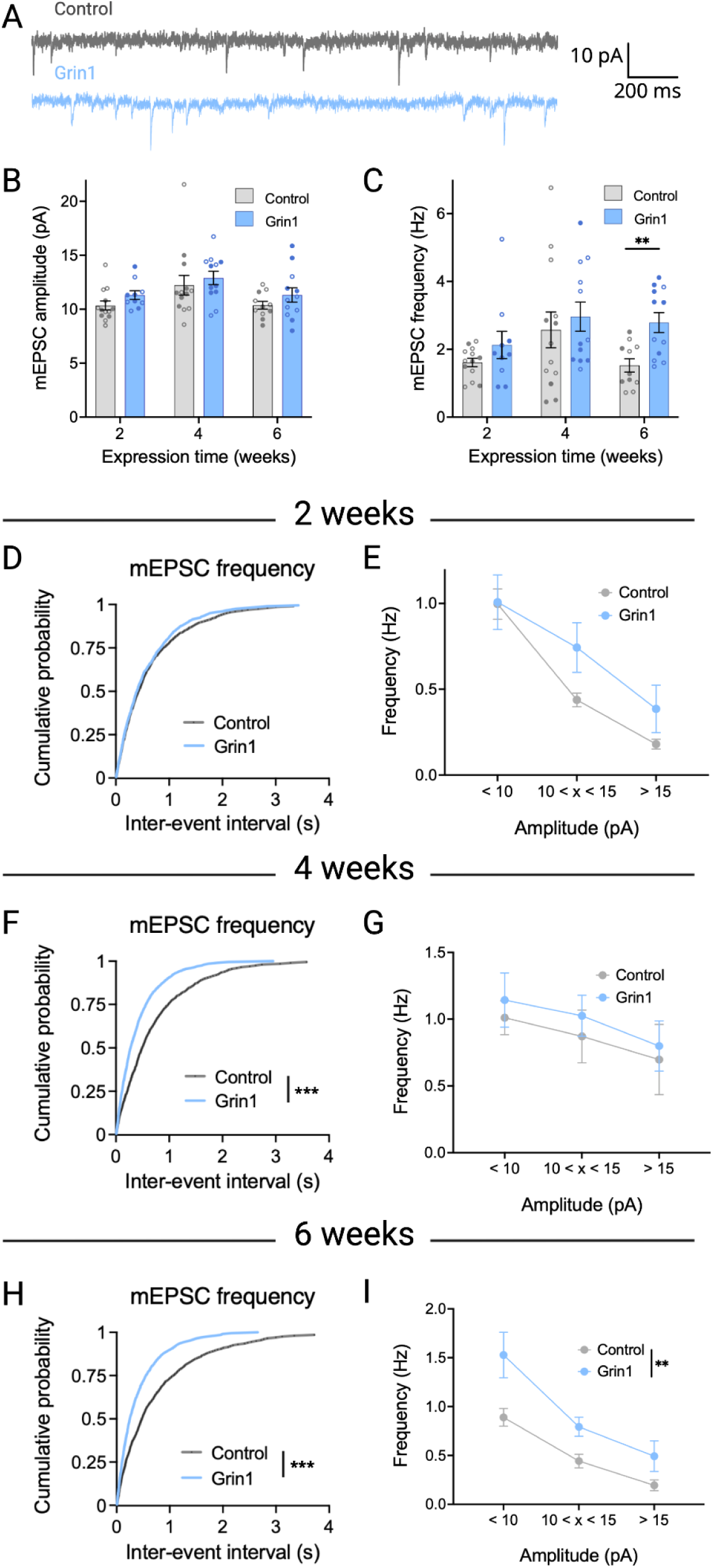
AMPAR-mediated synaptic transmission increases over time following NMDAR ablation. **(A)** Sample traces of miniature excitatory postsynaptic currents (mEPSCs) at 6 weeks. **(B)** No change in mEPSC amplitude at any time point (control n/N = 13/5, 13/6, and 11/5 at 2, 4 and 6 weeks; Grin1 n/N = 10/4, 12/4, and 12/6 at 2, 4 and 6 weeks). **(C)** mEPSC frequency is significantly increased at six weeks (main effect of virus: F_1,59_ = 6.063, P = 0.0167). **(D-E)** No change in the distribution of mEPSC frequency across all event sizes at two weeks. **(F)** There is a significant leftward shift in the distribution of mEPSC inter-event interval in the Grin1 group at four weeks (D = 0.2100, P = 0.0003) and **(H)** six weeks (D = 0.2150, P = 0.0002), with a similar effect across event amplitudes (main effect of virus at six weeks, F_1,21_= 12.12, P = 0.0022) **(G, I).** Sample sizes for panels C-I are the same as panel B. Data are mean ± s.e.m. for all panels; open and closed circles indicate recordings from female and male mice, respectively. **P < 0.01, ***P < 0.001; three-way ANOVA followed by simple effect test (B-C); Kolmogorov-Smirnov test (D, F, H); or two-way ANOVA (E, G, I); see Data S1 for complete statistics. https://BioRender.com/iazf88r

We also evaluated potential changes in mEPSC frequency as a function of event size. mEPSC amplitude can be influenced by synaptic location – i.e., events that originate at more distal synapses tend to have a smaller amplitude due to dendritic filtering, whereas events that originate closer to the soma tend to be larger [53]. At all timepoints, the relative difference in mEPSC frequency between Grin1 and control groups was similar, regardless of amplitude (Fig 3E, 3G, 3I).

To determine whether changes in spine density and excitatory synaptic transmission were caused by a cell-autonomous effect in pyramidal neurons, we used the same CRISPR-Cas9 genome editing strategy, but replaced the hSyn promoter sequence of our viral vectors with the CaMKII promoter sequence (Fig S2A). The efficacy of this manipulation was confirmed by a significant decrease in the NMDAR/AMPAR ratio 2 weeks after virus injection (Fig S2B-D). However, we did not observe significant changes in basilar spine density at this time point (Fig S2E-H). There were also no changes in mEPSC amplitude or frequency at any time point after Grin1 ablation using the CaMKII promoter (Fig S3), suggesting loss of NMDAR function from pyramidal neurons is not sufficient to cause the effects observed using the hSyn promoter.

### Compensatory Changes in Synaptic Transmission

Taken together, the changes in synaptic structure and function described above point towards an initial loss of dendritic spines in the Grin1 group. This is followed by a recovery of dendritic spine density that returns to and overshoots control levels, with a corresponding increase in mEPSC frequency. These changes may emerge as a compensatory response to the ablation of NMDARs, so we next investigated other potential compensatory changes in excitatory or inhibitory synaptic transmission. Following six weeks of virus expression, there was no change in the paired-pulse ratio of evoked EPSCs at interstimulus intervals ranging from 25-400 ms (Fig 4A-B). This indicates that presynaptic glutamate release probability was unaffected by NMDAR ablation, and also suggests the increase in mEPSC frequency at the same time point was related to dendritic spine density and synapse number, rather than a change in the probability of glutamate release.

**Figure 4.**
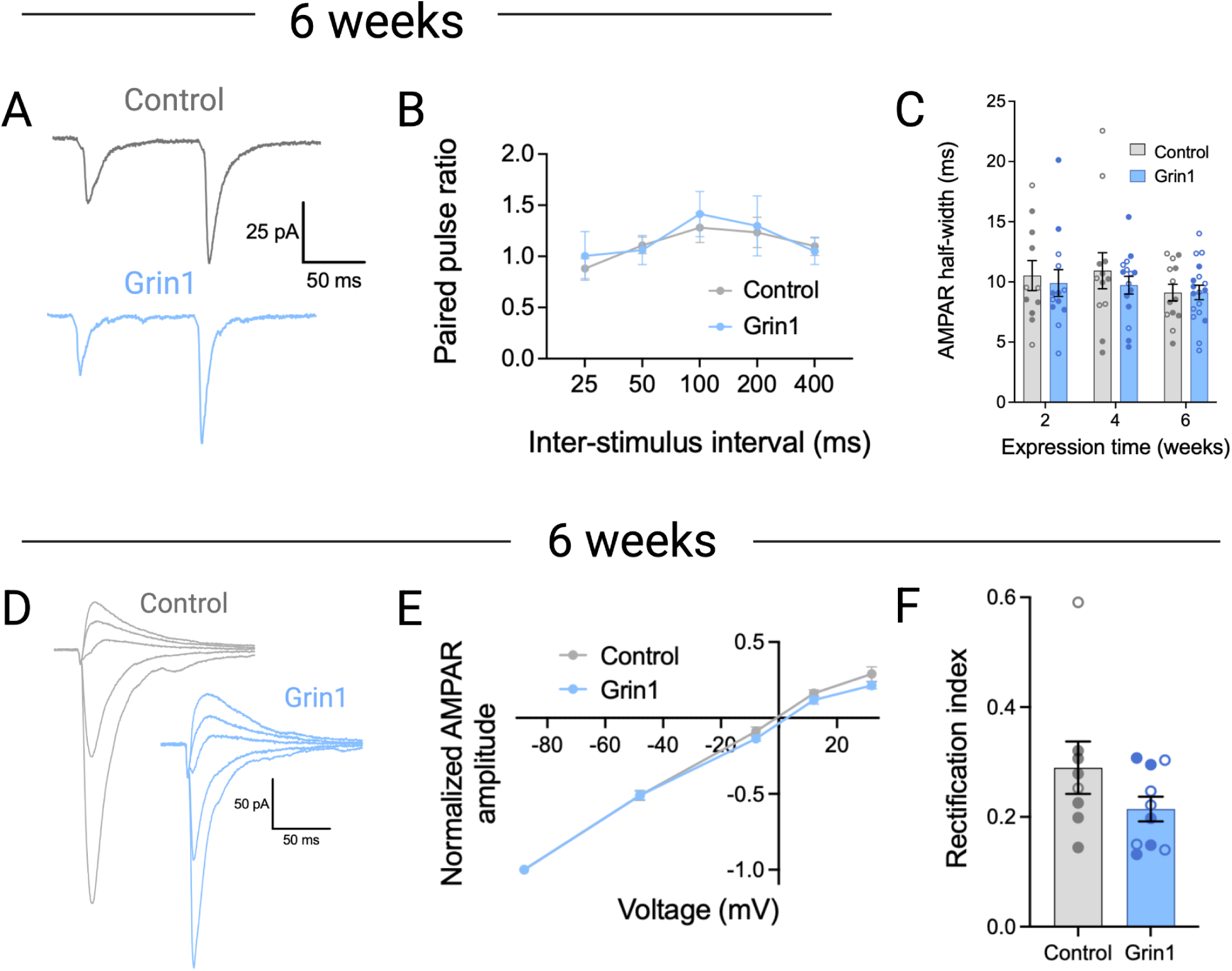
Paired-pulse ratio and AMPAR input-output properties are unchanged after NMDAR ablation. **(A)** Sample traces of EPSCs evoked in response to paired pulse stimulation after 6 weeks of virus expression. **(B)** No significant difference in paired pulse ratio (control n/N = 13-15/7; Grin1 n/N = 11-14/8). **(C)** Half width of the AMPA EPSC is unchanged at all time points (control n/N = 11/4, 12/5, and 13/6 at 2, 4 and 6 weeks; Grin1 n/N = 13/4, 15/6, and 18/6 at 2, 4 and 6 weeks). **(D)** Sample traces of AMPAR EPSCs evoked at different voltages. **(E)** AMPA current-voltage plot shows no significant difference between groups (control n/N = 6-12/7; Grin1 n/N = 3-13/7). **(F)** The rectification index is not significantly different between groups (control n/N = 8/4; Grin1 n/N = 10/7). Data are mean ± s.e.m. for all panels; open and closed circles indicate recordings from female and male mice, respectively. Mixed-effects analysis (B, E); three-way ANOVA (C); or Welch’s two-tailed t test (F); see Data S1 for complete statistics. https://BioRender.com/mlt9v8e

We next examined the possibility that NMDAR ablation and corresponding loss of calcium influx could lead to compensatory changes in AMPAR subunit composition, such as increased contribution of AMPARs lacking the GluA2 subunit [54,55]. Subunit composition can influence the kinetic properties of AMPAR currents, but there was no change at any time point in the half-width of the isolated AMPAR current recorded at −70 mV as part of the NMDAR/AMPAR ratio measurement (Fig 4C). Calcium-permeable AMPARs lacking a GluA2 subunit are blocked by intracellular polyamines at depolarized membrane potentials, producing a non-linear current-voltage relationship. However, following six weeks of virus expression, there was no difference in the AMPAR current-voltage relationship (Fig 4D-E), and no change in the rectification index comparing current passed at depolarized versus hyperpolarized membrane potentials (Fig. 4F). These data suggest that the subunit composition of AMPARs does not change following NMDAR ablation.

Given the increase in dendritic spine density and mEPSC frequency following six weeks of virus expression, we also explored potential adaptations at the level of inhibitory synaptic transmission, by recording miniature inhibitory postsynaptic currents (mIPSCs) at this time point (Fig 5A). Following six weeks of virus expression, we observed a significant increase in average mIPSC amplitude (Fig 5B-C), but no change in mIPSC frequency (Fig 5D-E). These results suggest that following protracted NMDAR ablation, the overall excitatory-inhibitory balance of the mPFC network may be maintained by stabilizing at a new allostatic set point (Fig. 5F-G).

**Figure 5.**
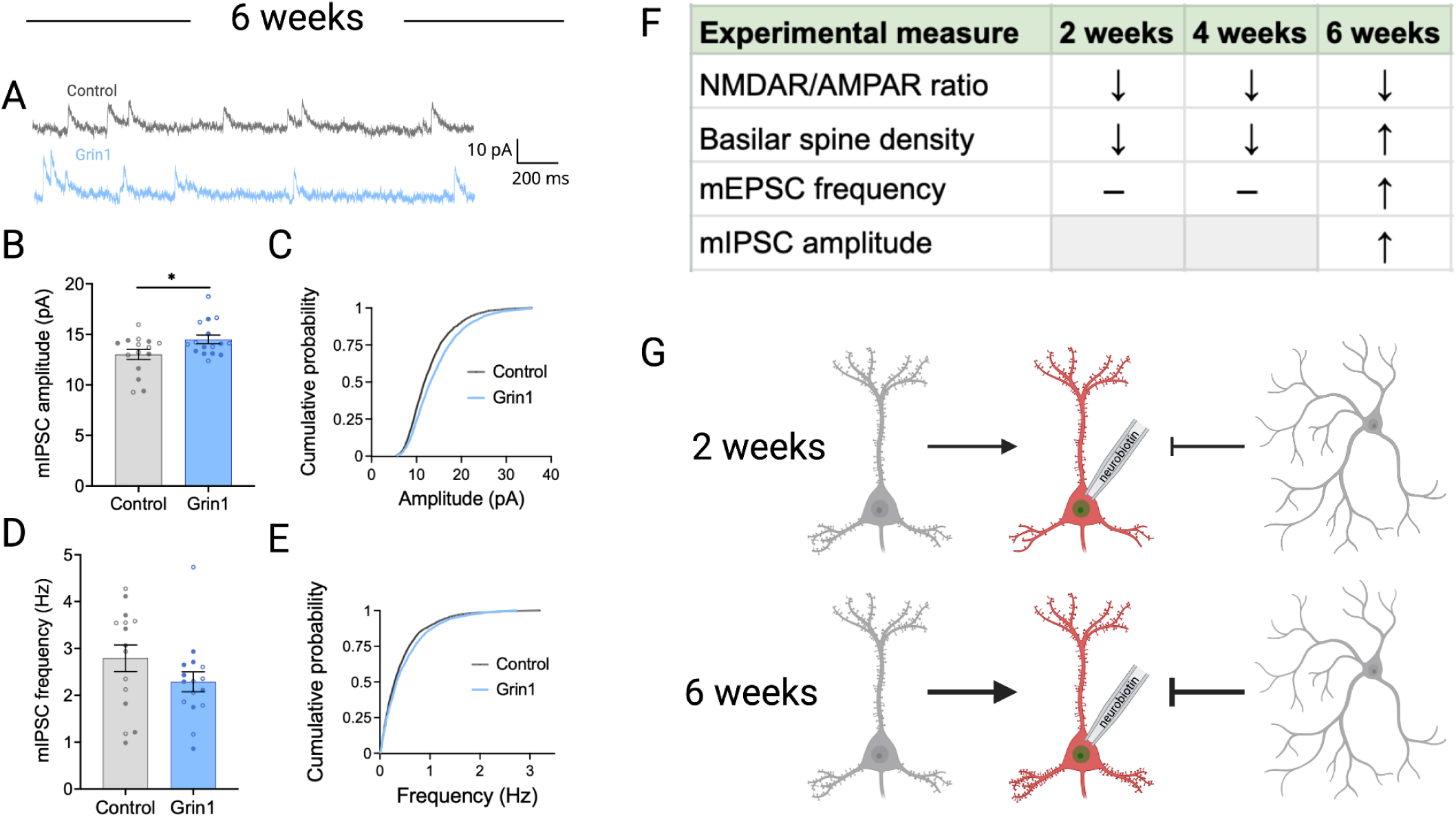
Inhibitory synaptic input to pyramidal neurons is increased after NMDAR ablation. **(A)** Sample traces of miniature inhibitory postsynaptic currents (mIPSCs) after 6 weeks of virus expression. **(B-C)** mIPSC amplitude is significantly increased in the Grin1 group (main effect of virus, F_1,27_ = 5.350, P = 0.0286), with no change in the cumulative probability distribution**. (D-E)** mIPSC amplitude is unchanged following NMDAR ablation (for B-E: control n/N = 15/5; Grin1 n/N = 16/4). Data are mean ± s.e.m. for all panels; open and closed circles indicate recordings from female and male mice, respectively. *P <0.05; two-way ANOVA (B, D); or Kolmogorov-Smirnov test (C, E); see Data S1 for complete statistics. **(F)** Summary table and figure **(G)** illustrating functional and structural changes over time following NMDAR ablation. https://BioRender.com/ilmaze5

## DISCUSSION

Our findings reveal significant time-dependent structural and functional reorganization of cortical networks following NMDAR manipulation during adolescence. We observed a change in basilar spine density as NMDAR ablation progressed, with initial spine loss in the Grin1 group at earlier time points shifting towards an increase in spine density at 6 weeks. This effect was primarily driven by thin spines, which are associated with weaker synapses and appear or disappear in response to synaptic activity [56], as well as filopodia, which have a short lifespan and may not form functional synapses due to their lack of postsynaptic density [52]. This is consistent with evidence from human postmortem tissue studies showing a selective loss of smaller spines in the cortex of patients with schizophrenia [57,58]. We did not observe a change in apical spine density at any time point. This suggests that the local mPFC microcircuitry may be more vulnerable to the effects of NMDAR ablation, since basilar dendrites receive inputs from other nearby neurons, whereas apical dendrites receive more long-range cortical and thalamic signals [59]. This basilar-specific effect is consistent with evidence of decreased synchronous firing between pairs of prefrontal neurons following NMDAR blockade [16], which indicates that disruption of local PFC connections may be a key consequence of NMDAR hypofunction.

We were surprised that changes in dendritic spine architecture were observed in the basilar but not the apical dendritic arbor of layer V pyramidal neurons. Studies of postmortem tissue from patients with schizophrenia have documented a reduction in dendritic spine density in the basilar dendrites of deep layer III neurons in the dorsolateral PFC (dlPFC). Based on these previous studies, it has been suggested that apical dendrites in layer V neurons may be vulnerable to spine loss in schizophrenia, since deep layer apical dendrites and superficial layer basilar dendrites both receive inputs from the mediodorsal nucleus of the thalamus within layer III [60]. In contrast, the significant increase in basilar spine density that we observed at 6 weeks suggests that the network is overcompensating for early initial spine loss. This may be attributed to the selective nature of our manipulation; while spine density decreased as we predicted at 2 and 4 weeks, neurons were able to adapt by producing more spines because we did not alter synaptic architecture proteins that are implicated in spine loss in schizophrenia, such as Cdc42 and Duo [60]. This would also explain why we observed an increase in spine density at later time points, despite our initial prediction that loss of NMDARs would lead to a reduction in synapse density due to activity-dependent disconnection [16]. Our findings indicate that NMDAR ablation alone is not sufficient to induce long-lasting deficits in spine density, and suggests that other risk factors for the disease – whether genetic or environmental – could also play a role in the failure to recover from loss of neuropil observed in patients with schizophrenia.

We examined whether the changes in spine density were associated with functional consequences at the synaptic level, and observed a significant increase in mEPSC frequency at 6 weeks. There were no compensatory change in presynaptic glutamate release or AMPAR subunit composition, indicating that the mEPSC frequency increase was primarily driven by the alterations in basilar spine density. Interestingly, we also observed an increase in inhibitory signaling, with mIPSC amplitude significantly greater in the Grin1 group at 6 weeks. It was surprising that there were no mEPSC changes observed at 2 weeks, given the significant reduction in basilar spine density at this early time point. This may be attributed to the fact that early spine loss was primarily due to a reduction in weaker, less stable spines such as thin spines and filopodia, which may make a smaller contribution to detectable levels of excitatory synaptic transmission [52]. In contrast, mushroom spine density has a much smaller decrease in Grin1 mice at 2 weeks. At 6 weeks, the density of mushroom spines, which are more stable and associated with stronger synapses [61], is equivalent in the Grin1 and control groups and the density of weaker spines is significantly greater, which may be sufficient to produce the observed increase in mEPSC frequency. Overall, these results suggest that subtle changes in synaptic architecture may precede functional consequences for the network, with initial loss of immature spines leading to a compensatory increase in connectivity and neurotransmission at later time points.

A key question in the field is whether pyramidal neurons or interneurons are primarily affected by NMDAR hypofunction. Abnormal glutamatergic postsynaptic proteins and excessive synaptic pruning in excitatory neurons [62–65] may lead to a subsequent downregulation of feedback inhibition, as reflected by decreased interneuron expression of the activity-dependent proteins glutamate decarboxylase 67 and parvalbumin [62,66,67]. However, we found that direct manipulation of NMDAR function in pyramidal neurons (using the CaMKII promoter) did not recapitulate the effects of pan-neuronal NMDAR ablation (using the hSyn promoter). This suggests that network reorganization may either require simultaneous loss of NMDAR function in both pyramidal neurons and inhibitory interneurons, or that loss of NMDAR function from inhibitory interneurons may instigate network reorganization. Several lines of evidence are consistent with the latter possibility. Patients with schizophrenia have lower GABA synthesis [68], and reduced expression of interneuron neuropeptides and GABA-A receptor subunits [69]. Reduced GAD67 expression is also sufficient to induce schizophrenia-related behavioral phenotypes and altered excitatory and inhibitory synaptic transmission [70,71]. In addition, interneurons are preferentially targeted by NMDAR antagonists due to their more depolarized membrane potential [72], raising the question of whether previous studies using NMDAR antagonism as a model for schizophrenia have exerted greater effects upon inhibitory neurons than excitatory neurons. These issues can be addressed in future research using genetic manipulations developed to target interneurons [73–77].

In addition to future studies targeting more specific cell types, our genome editing platform can be readily adapted to manipulate expression of other genes strongly associated with schizophrenia, simply by changing the viral vector gRNA sequence. These could include the Grin2a gene that encodes the GluN2A subunit of NMDARs, or the Gria3 gene that encodes the GluA3 subunit of AMPA receptors [78]. While schizophrenia is unlikely to involve complete loss of NMDAR function (e.g., our 6 week timepoint), the partial loss of NMDAR function we observe at earlier timepoints may have greater translational relevance. The progressive nature of our manipulation is thus valuable for correlating degree of NMDAR disruption with network reorganization and behavioral output across time. In a parallel study, we have used this strategy to track cognitive impairments produced by mPFC NMDAR ablation in a restless bandit task [79]. These cognitive impairments develop within two weeks, and recover partially but not fully over time. Behavioral deficits may persist because upregulation of AMPAR function cannot fully restore network operations normally mediated by the unique biophysical properties of NMDARs, such as persistent neuronal firing during working memory [80].

Our data address questions regarding the emergence of compensatory synaptic mechanisms, which are central to developing our understanding of schizophrenia. Symptom onset may be triggered in patients by the interaction of multiple layered risk factors [81]. One theory of the disease is that genetic mutations affecting synapse formation and maintenance cause latent defects in cortical connectivity that are masked by the hyperconnected state of the brain in early development, but as extraneous synapses are pruned during adolescence, these vulnerabilities are laid bare and symptoms begin to occur in patients [38] In our model, the cellular mechanisms of spine formation and maintenance were unchanged, and therefore early synaptic defects could be compensated for by the production of additional immature spines. Although disruptions to the Grin1 gene alone cannot fully recapitulate the pathophysiology of schizophrenia, studying the effects of an isolated Grin1 manipulation may help us understand why first-degree relatives of patients who have Grin1 mutations but no additional genetic risk factors do not develop schizophrenia, or why patients with fewer biological or environmental risk factors are able to successfully recover from first-episode psychosis. In contrast, other patients may have altered gene expression related to neuronal survival [82] and architecture, such as complement-mediated synaptic pruning [64,83], thus limiting the brain’s ability to adapt to insults and leading to more severe or long-lasting symptoms. Further study will be needed to examine how synaptic changes such as NMDAR ablation interact with other risk factors to influence processes of synaptic dysfunction and compensation relevant to schizophrenia. However, the preclinical success of newly developed therapies to treat NMDAR hypofunction is an encouraging sign that these synaptic deficits and associated cognitive symptoms may be able to be restored in patients [84].

## ACKNOWLEDGEMENTS

Viral vectors used in this study were generated by the University of Minnesota Viral Innovation Core and the Center for Neural Circuits in Addiction (P30 DA048742). We thank the University of Minnesota University Imaging Core (SCR_020997) for the use of their facilities to conduct confocal imaging and analysis of dendritic spines, and Dr. Matthew Hearing for advice regarding electrophysiology experiments.

## AUTHOR CONTRIBUTIONS

RMD, NMG, and PER were responsible for study design. RMD, APA, and CRQ conducted and analyzed electrophysiology recordings. RMD performed immunohistochemistry protocols. RMD and HA performed intracranial injections of viral vectors generated by EMFV. RMD, LBC, ATP, HA, and JSM conducted confocal imaging and analysis of dendritic spines. RMD and PER wrote the original draft of the manuscript, which all authors reviewed and edited before submission.

## FUNDING

This work was supported by grants from the National Institutes of Health (T32MH115886 and F31MH133285 to RMD; T32GM008244, T32DA007234, and F30 DA060027 to APA; P50MH119569 to NHG and PER); the Howard Hughes Medical Institute (GT17865 to CRQ); the University of Minnesota’s MnDRIVE (Minnesota’s Discovery, Research, and Innovation Economy) initiative (PER); the University of Minnesota Life Sciences Summer Undergraduate Research Program (ATP); and the University of Minnesota Undergraduate Career Opportunities in Neuroscience and Undergraduate Research Opportunities Program (HA).

## COMPETING INTERESTS

The authors declare no competing interests.

## DATA AVAILABILITY STATEMENT

Data are available from the corresponding author upon request.

## Notes

### Competing Interest Statement

The authors have declared no competing interest.

### Summary of Updates

Revised text and new supplementary data

