## Supplemental Material for "Synaptic Dysfunction and Compensation After NMDA Receptor Ablation in the Mouse Medial Prefrontal Cortex"

**SUPPLEMENTARY MATERIALS & METHODS**

**Animals**

Mice were housed on a 12-h light cycle (0600–1800 h) at ~23ºC with ad libitum access to food and water. All experimental procedures were conducted between 0800h – 1800h, using comparable numbers of female and male mice in each experiment.

**Surgical Procedures**

Adeno-associated viral vectors were generated by the Viral Innovation Core at the University of Minnesota. We targeted the Grin1 gene for genome editing using a guide RNA (gRNA) sequence [CCGCGCCGATGTTGACAATCT; "gRNA(Grin1)"] expressed using AAV9-U6-gRNA(Grin1)-hSyn-mCherry-Cre (titer: 2.16E13 GC/ml), with a control gRNA targeting LacZ [TGCGAATACGCCCACGCGAT; "gRNA(LacZ")] expressed using AAV9-U6-gRNA(LacZ)-hSyn-mCherry-Cre (titer: 1.75E13 GC/ml) [44]. Both viral vectors expressed gRNA in tandem with an mCherry-Cre fusion protein to visualize transduced cells and catalyze Cre-dependent Cas9 expression for genome editing, under the control of the human synapsin (hSyn) promoter to target all neurons. We constructed additional viral vectors in which the hSyn promoter was replaced by a 0.4 kb α-calcium/calmodulin-dependent protein kinase II (CaMKII) promoter [1], to more selectively target excitatory projection neurons.

Mice were anesthetized with a ketamine/xylazine cocktail (100/10 mg/kg, i.p.), and surgical procedures were performed using a stereotaxic alignment system (Kopf Instruments, Tujunga, CA, USA) with a heating pad to maintain body temperature. After infusion, the needle was left in place for five minutes to allow for viral diffusion, then gradually retracted over the course of approximately two minutes. Mice were administered carprofen (10 mg/kg, s.c.) on the day of surgery and for three subsequent days.

**Immunohistochemistry for virus expression**

Coronal mPFC sections were cut in PBS at 50 microns on a vibratome (Leica VT1000S) and stored at -20ºC in a cryoprotectant solution containing 50 mM phosphate buffer, 876 mM sucrose, and 30% ethylene glycol. Slices were washed 3 x 15 mins in PBS, incubated with blocking buffer (PBS with 2% normal horse serum (NHS) and 1% Triton-X100) for two hours at room temperature, then incubated with chicken polyclonal anti-GFP (Abcam ab13970, 1:2000) for 72 hours at 4ºC. We did not include a primary antibody for mCherry because the endogenous mCherry signal was bright enough to be detected without amplification. Sections were washed 3 x 15 minutes with PBS and incubated overnight with donkey anti-chicken Alexa Fluor 488 (Jackson Immuno Research Labs item 703-545-155, 1:300) at 4ºC. Slices were washed once for 15 minutes with PBS, incubated with DAPI nucleic acid stain (Invitrogen, 1:5000) in PBS with 0.03% Triton-X100, 1% bovine serum albumin, 1% normal donkey serum, and 0.01% sodium azide for 20 min, and washed three times with PBS for 15 min, 90 min, and 120 min. After staining, slides were imaged on a Keyence BZX fluorescence microscope under a 10x objective using mCherry and DAPI filter cubes. BZX Analyzer software was used to stitch multi-channel tiled 10x z-stack images into a single image.

**Brain slice electrophysiology**

Mice were deeply anesthetized with isoflurane and perfused with 10 mL ice-cold sucrose cutting solution containing (in mM): 200 sucrose, 1.9 KCl, 1.2 $NaH_2PO_4$, 33 $NaHCO_3$, 10 glucose, 0.4 ascorbic acid, 6 $MgCl_2$, 0.5 $CaCl_2$. Mice were subsequently decapitated and brains were quickly removed then placed in ice-cold sucrose cutting solution. Coronal slices (240 μm thick) containing mPFC were collected using a vibratome (Leica VT1000S) and allowed to recover while submerged in artificial cerebrospinal fluid (aCSF) containing (in mM): 119 NaCl, 2.5 KCl, 1 $NaH_2PO_4$, 26.2 $NaHCO_3$, 11 glucose, 0.4 ascorbic acid, 4 $MgCl_2$, 1 $CaCl_2$. Slices recovered in warm aCSF (33ºC) for 10 min and then equilibrated to room temperature for at least thirty minutes before use. Slices were transferred to a submerged recording chamber and continuously perfused with recording aCSF containing (in mM): 125 NaCl, 2.5 KCl, 25 $NaHCO_3$, 10 glucose, 0.4 ascorbic acid, 1.3 $MgCl_2$, 2 $CaCl_2$ at a rate of 2 mL/min at room temperature. All solutions were continuously oxygenated (95% $O_2$/5% $CO_2$).

Virally transduced pyramidal neurons in layer V of the mPFC (including anterior cingulate, prelimbic, and infralimbic cortex) were identified by expression of mCherry using an Olympus BX51W1 microscope. Whole-cell voltage-clamp recordings were made with borosilicate glass electrodes (3–4.5 MΩ) filled with (in mM): 120 $CsMeSO_4$, 15 CsCl, 10 TEA-Cl, 8 NaCl, 10 HEPES, 5 EGTA, 5 QX-314, 4 ATP-Mg, 0.3 GTP-Na and 0.13% neurobiotin (pH 7.2-7.3). For AMPAR current-voltage measurements, spermine (0.1 mM) was added to the internal solution, with correction for a liquid junction potential of approximately 8 mV in this experiment only.

Excitatory postsynaptic currents (EPSCs) were electrically evoked at 0.33 Hz using a glass monopolar electrode placed in layer V/VI, and pharmacologically isolated using GABA-A receptor antagonist picrotoxin (50 μM, Tocris) and GABA-B receptor antagonist SCH 59011 (10 μM, Tocris). AMPAR-mediated EPSCS were measured at a holding potential of -70 mV. NMDAR-mediated EPSCs were measured 50 ms following stimulation at a holding potential of +40 mV. Inhibitory postsynaptic currents (IPSCs) were measured at +10 mV and pharmacologically isolated using the NMDAR antagonist D-APV (50 μM, R&D Systems) and the AMPAR antagonist NBQX (10 μM, R&D Systems). Miniature synaptic currents (mEPSCs and mIPSCs) were measured in the presence of tetrodotoxin (500 nM, Fisher Scientific) to block the generation of action potentials.

All recordings were performed using a MultiClamp 700B amplifier (Molecular Devices) and Dendrite digitizer (Sutter Instrument), filtered at 2 kHz, and digitized at 10 kHz. Data acquisition was performed using SutterPatch software version 2.3 in Igor Pro version 8.04 (WaveMetrics). Easy Electrophysiology software version 2.8.0 (RRID:SCR_021190) was used for analysis of synaptic currents. For mEPSCs and mIPSCs, at least 200 events per cell were acquired across 20 second sweeps, filtered at 1.5 kHz and detected using an amplitude threshold of 5 pA and a detection criterion of 4.00 [2].

**Confocal imaging and analysis of dendritic spines**

Brain slices containing neurons filled with neurobiotin were fixed in 4% PFA for a minimum of 12 hours, washed 3 x 15 minutes with PBS, and blocked with a solution containing 2% NHS in PBS with 0.05% Tween-20 and 0.2% Triton-X100 for 3 hours at room temperature. Slices were incubated for 18 hours in Cy5-streptavidin (Cytiva – Amersham) diluted 1:500 in the same blocking solution at 4ºC. Sections were rinsed 3 x 15 minutes in PBS containing 0.1% Tween-20, mounted from DI $H_2O$, and coverslipped onto glass slides using ProLong Gold Antifade with DAPI mountant (Thermo Fisher Scientific).

For confocal imaging, the laser's excitation wavelength was set to 644 nm to image Cy5-streptavidin while minimizing signal from viral mCherry expression. Cells were imaged first with a 20x oil immersion objective 0.75 NA with an xy pixel size of 568 nm and a z-step size of 0.96 microns to provide an overall view of the dendritic arbor, and then spines were imaged with a 63x oil immersion objective 1.4 NA with an xy pixel size of 93 nm and a z-step size of 0.28 microns. Apical dendrites of layer V neurons were imaged in layer I of the mPFC, which is the site of synaptic contacts between layer V and layer II/III mPFC neurons [3]. The layer V neuron's longest basilar dendrite and its branches were also imaged [4,5]. Lightning deconvolution was applied to all 63x images to improve resolution. For spine analysis in Imaris, images were first masked to remove background signal, and filaments were manually drawn to capture the path of the dendrites. Spines were detected using the seed point classification algorithm and manually corrected to remove duplicate seed points and/or add missing spines as needed. Spine morphology was automatically classified using the built-in Matlab toolbox in Imaris.

**SUPPLEMENTARY FIGURES & FIGURE LEGENDS**

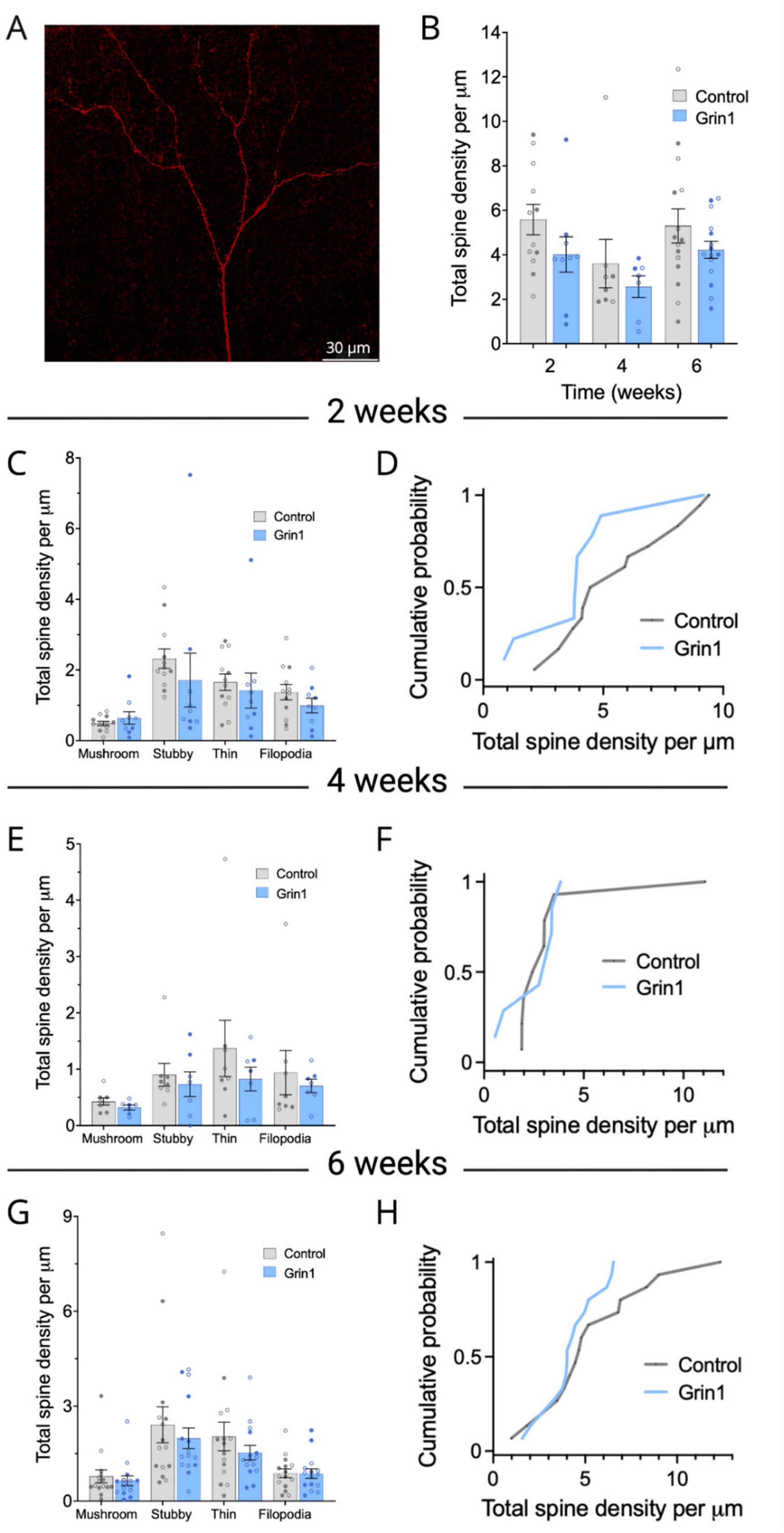


**Figure S1. Apical spine density is unchanged following NMDAR ablation. (A)** Sample image of the apical arbor of a layer V mPFC pyramidal neuron at 63x magnification. **(B)** No significant changes in total apical spine density in the Grin1 group at any timepoint (control n/N = 12/5, 8/6, and 15/8 for 2, 4, and 6 weeks; Grin1 n/N = 9/4, 7/4, and 15/9 for 2, 4, and 6 weeks). No significant change in apical spine density across all morphological categories and no difference in the cumulative probability distribution at two weeks **(C-D)**, four weeks **(E-F)**, or six weeks **(G-H)** (all sample sizes same as B). Data are mean ± s.e.m. for all panels; open and closed circles indicate recordings from female and male mice, respectively. Three-way ANOVA (B); two-way ANOVA (C, E, G); or Kolmogorov-Smirnov test (D, F, H); see Data S1 for complete statistics.

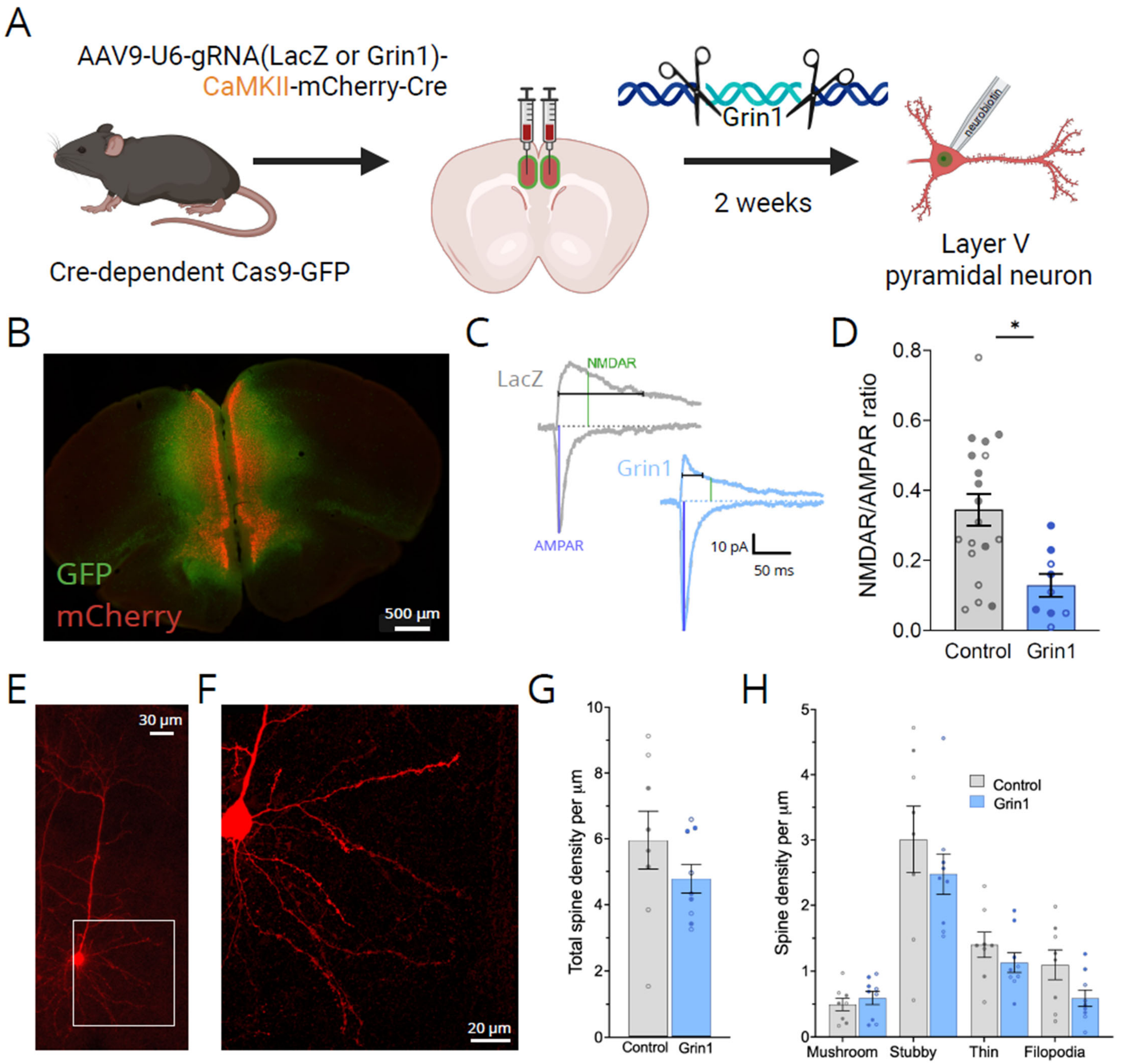


**Figure S2.** **(A)** Experimental overview showing mPFC injection of CaMKII-driven virus to ablate Grin1 gene expression, followed by whole-cell patch-clamp recording and neurobiotin cell filling. **(B)** Expression of GFP and mCherry following virus injection into mPFC (10x magnification tiled image). **(C)** Sample traces of EPSCs at -70 mV and +40 mV to measure AMPAR and NMDAR currents, respectively. **(D)** Grin1 ablation significantly reduces NMDAR/AMPAR ratio after 2 weeks of virus expression, with a significant main effect of Virus ($F_{1,24}$ = 7.69, p = 0.011) (n/N=19/6 and 9/4 for LacZ and Grin1, respectively). **(E)** Sample image of a neurobiotin-filled layer V mPFC pyramidal neuron at 20x magnification (dendrites of neighboring cell also shown). **(F)** Sample image of the basilar arbor of the same neuron at 63x magnification. There is no significant change in total basilar spine density **(G)** or across any individual morphological category **(H)** (control n/N = 8/6, Grin1 n/N = 9/5). Data are mean ± s.e.m. for all panels; open and closed circles indicate recordings from female and male mice, respectively. *p < 0.05, ANOVA main effect of Virus; see Data S1 for complete statistics.

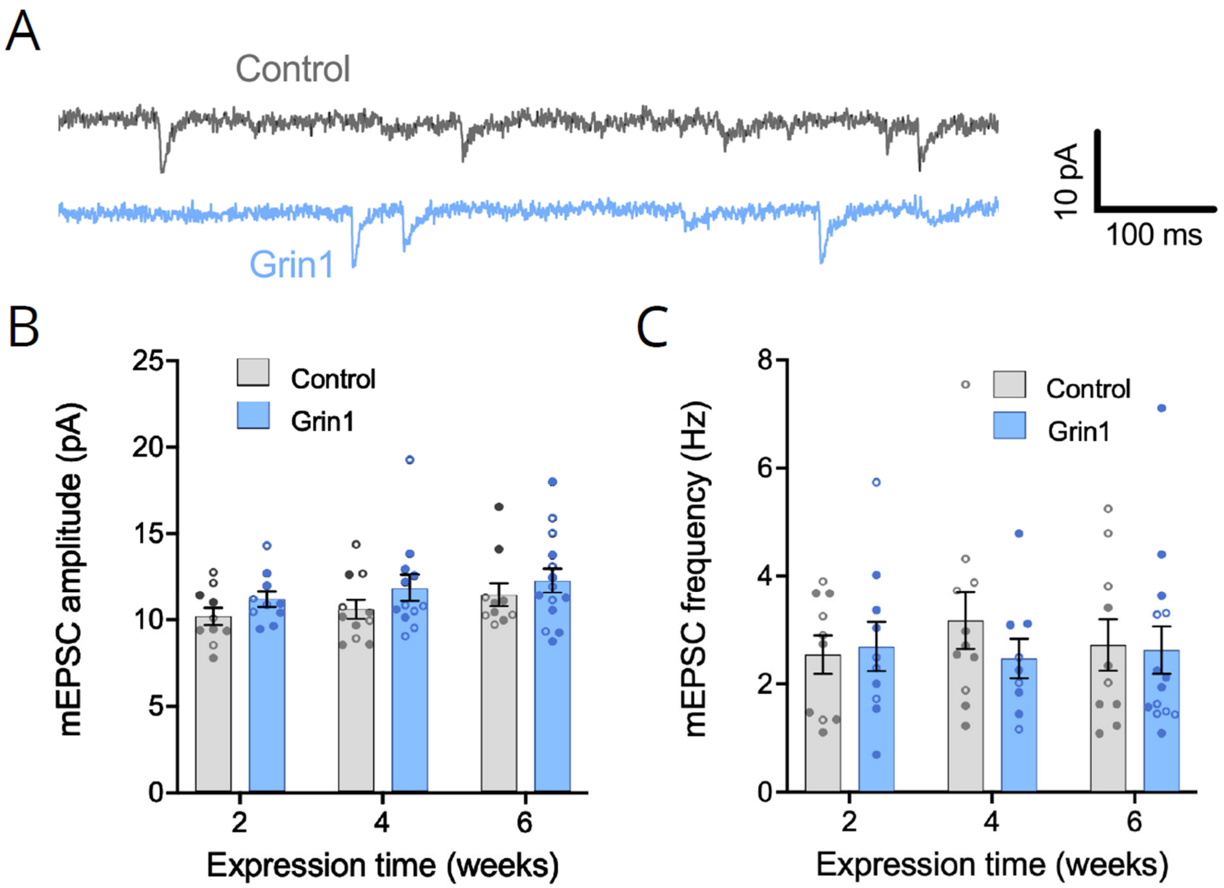


**Figure S3**. Grin1 ablation driven by the CaMKII promoter does not change mEPSC amplitude or frequency. **(A)** Sample traces of miniature excitatory postsynaptic currents (mEPSCs) after 6 weeks of control or Grin1 virus expression. There were no significant changes in mEPSC amplitude **(B)** or frequency **(C)** at any time point (control n/N = 10/6, 11/5, and 10/4 at 2, 4, and 6 weeks; Grin1 n/N = 10/6, 9/5, and 14/6 at 2, 4, and 6 weeks). Data are mean ± s.e.m. for all panels; open and closed circles indicate recordings from female and male mice, respectively. Three-way ANOVA (B,C); see Data S1 for complete statistics.
